# Cell size matters: nano- and micro-plastics preferentially drive declines of large marine phytoplankton due to co-aggregation

**DOI:** 10.1101/2021.08.31.458382

**Authors:** Craig J. Dedman, Joseph A. Christie-Oleza, Víctor Fernández-Juárez, Pedro Echeveste

## Abstract

Marine plastic pollution represents a key environmental concern. Whilst ecotoxicological data for plastic is increasingly available, its impact upon marine phytoplankton remains unclear. Owing to their predicted abundance in the marine environment and likely interactions with phytoplankton, here we focus on the smaller fraction of plastic particles (∼50 nm and ∼2 µm polystyrene spheres). Exposure of natural phytoplankton communities and laboratory cultures revealed that plastic exposure does not follow traditional trends in ecotoxicological research, since large phytoplankton appear particularly susceptible towards plastics exposure despite their higher surface-to-volume ratios. Cell declines appear driven by hetero-aggregation and co-sedimentation of cells with plastic particles, recorded visually and demonstrated using confocal microscopy. As a consequence, plastic exposure also caused disruption to photosynthetic functioning, as determined by both photosynthetic efficiency and high throughput proteomics. Negative effects upon phytoplankton are recorded at concentrations orders of magnitude above those estimated in the environment. Hence, it is likely that impacts of NPs and MPs are exacerbated at the high concentrations typically used in ecotoxicological research (*i*.*e*., mg L^-1^).

## Introduction

The occurrence of plastic debris within the global ocean appears ubiquitous and represents a major environmental concern.^1, 2^ Estimates predict that 4.8 to 12.7 million tons of plastic enter the marine environment every year,^3^ and currently up to 5.35 trillion plastic particles are believed to be carried along ocean currents where they may interact with marine species.^4^ Research pertaining to the environmental fate of plastic pollution and its interaction with biota has increased dramatically over the past decade.^1, 2, 5, 6^ Here, investigation has primarily focussed upon the occurrence of plastic particles <5 mm, termed microplastics.^1, 2, 7^ Such particles have been found to display the potential to exert adverse impacts upon marine biota spanning various trophic levels and may ultimately affect human health through the consumption of marine aquaculture products.^1, 8-10^

Microplastics are believed likely to continually fragment in the natural environment until completely mineralised, forming particles of an ever-decreasing size, eventually giving rise to nanoplastics (<1 µm).^2, 11-14^ This definition is still under debate, where others define nanoplastics as those particles <100 nm, in-line with the classification of metal- and metal oxide nanomaterials.^2, 11-16^ The fragmentation of microplastics into nanoplastics has been demonstrated under laboratory conditions.^11, 17^ For example, shear forces which occur during wastewater processing is observed to cause fragmentation of microplastics released from ‘facial scrubbers’ found in facewash.^17^ It is believed that in the environment, one microplastic particle (5 mm) could fragment into 10^14^ nanoplastics sized 100 nm.^14, 18, 19^ As such, the quantity of nano-sized plastic debris in the environment is likely to far exceed those predicted for microplastics.

Environmental sampling highlights a missing fraction of plastic particles <1 mm, believed the cause of underestimations of the global oceanic load of plastic debris.^6, 20^ As such, uncertainties still exist in the true environmental concentration of small plastic particles. The abundance and distribution of plastic particles sized <250 µm across the global ocean is not known.^21^ Analytical techniques have been insufficient to accurately ascertain concentrations of these small plastic particles in the environment, which often pass through mesh during sampling.^5, 18, 21-23^ Hence, researchers face great difficulty in correctly assessing the plastic budget of particles below the µm range. However, recent field investigations carried out by Pabortsava and Lampitt (2020) revealed that small plastic fragments which are contained within the ocean interior can account for the “missing sink” of plastic previously described. Based on the evidence currently available the concentration of small plastics in the µm and nm size range appears to be in the low µg L^-1^ range.^21, 24^ Novel approaches are being developed to accurately monitor and characterise small plastic fragments, requiring multiple analytical techniques working simultaneously.^22^ Despite analytical challenges, nanoplastics have been identified in the environment.^25^ For instance, Ter halle *et al*. (2017) reported the occurrence of various polymers of nanoplastics within the North Atlantic subtropical gyre. Uncertainty regarding the true volume of plastic pollution within the global ocean is likely to change as our analytical capabilities advance. Due to differing physical and chemical behaviour in the nanoscale, nanoplastics are likely to display significantly altered fate within the marine environment compared to microplastics.^18, 26, 27^ Their small size increases the surface area-volume ratio of particles, increasing reactivity and enhancing their ability to absorb other contaminants within the water column, as well as enhancing bioavailability.^13, 18^

The marine phototrophic community contributes approximately 50% of global primary productivity,^28^ and contributes to major biogeochemical and climatic cycles, as well as occupying the base of the marine food web.^28^ Whilst the impact of plastic particles upon biota has increasingly been studied, effects upon marine phytoplankton remain relatively little understood.^29, 30^ These organisms are often the first to interact with contaminants of anthropogenic origin and are crucial to maintaining the health of the marine ecosystem. Plastic exposure has previously been associated with a reduction in fitness of marine phytoplankton,^29^ often characterised by reductions in algal growth and photosynthetic performance.^29, 31, 32^ Adverse effects of plastics exposure have been recorded in green algae,^32-34^ diatoms^31, 34-38^ and cyanobacteria.^39-41^ However, findings vary,^42-44^ and, in particular, evidence for the potential impact of nanoplastics upon marine phytoplankton is limited.^18, 34, 45^ While this is the case, preliminary evidence suggests that nano-sized plastic particles may be more detrimental to phytoplankton their larger counterparts.^31, 44, 46^ The ocean represents a significant carbon sink and any adverse impact of plastic exposure could result in damages to oceanic carbon sequestration,^30^ a major concern given the threat of climate change. As such it is of high importance that additional research is carried out to reveal the possible impact of plastic particles spanning the micron-nanometre size range upon this ecologically important group of species.

Herein, we investigate the impact of ∼2 µm polystyrene (PS) microspheres (MPs) and ∼50 nm polystyrene nanoparticles (NPs), representing the small plastic debris predicted to be abundant in the environment, upon marine phytoplankton. As mentioned above, the exact concentrations of such plastic debris in the environment remain uncertain,^47^ hence in our work supra-environmental concentrations (∼100-100,000X those predicted in the environment) are utilised to provide insight into potential effects of exposure and to identify any threshold values for toxicity.^47, 48^ First, we examined the natural marine community response towards plastics exposure, revealing a clear relationship between cell size and detrimental impacts upon phytoplankton growth. Here, greatest adverse effects were observed on larger eukaryotic species, whilst picocyanobacteria were little affected. This trend was subsequently confirmed by exposure of a range of laboratory cultured phytoplankton taxa varying in cell size and taxonomy in natural seawater. Adverse impacts were only recorded at concentrations exceeding those predicted in the environment. Using *Emiliania huxleyi* as a model, hetero-aggregation was revealed to be the key mechanism facilitating the decline of large phytoplankton in the presence of small plastic particles. Entrapment of cells within clusters of plastic aggregates and subsequent removal from the water column, alongside disruption to photosynthetic processes appeared the key features of plastic toxicity at supra-environmental concentrations.

## 2. Methods

### 2.1 Materials

Plastic particles used during experimental work were obtained from Spherotech™. Nanoplastics (NPs) and microplastics (MPs) were sized approximately 50 nm and 2 µm, respectively (see individual sections for specific details of particles used). Reagents used during experiments were obtained from Sigma Aldrich. Natural seawater (NSW) routinely used in experimental work was collected from Station L4, Plymouth (50°15.0’N; 4°13.0’W) and autoclaved prior to use.

### 2.2 Phytoplankton cultures

Axenic phytoplankton cultures were routinely grown onsite at the University of Warwick: cyanobacteria *Prochlorococcus sp*. MED4 and *Synechococcus sp*. WH7803, green algae *Micromonas sp*. CMP2709 and *Ostreococcus tauri* OTH95, haptophyte *Emiliania huxleyi* CCMP1516, and diatoms *Thalassiosira pseudonana* CCMP1335 and *Phaeodactylum tricornutum* CCMP2561. Before plastic exposure, the phototrophs were routinely grown in axenic enriched media, ideal for each species as suggested by the literature (artificial seawater for *Synechococcus*;^49^ Pro99 for *Prochlorococcus*;^50^ K-media for green algae and *E. huxleyi*;^51, 52^ and F/2 media for diatoms).^53^

### 2.3 Monitoring the microbial community response towards micro- and nano-plastic exposure using flow cytometry

Natural seawater (NSW) and its associated microbial community was collected from a pristine coastal location in Mallorca (39.493868 - 2.739820, November 2020) at a depth of 1 m and returned to the laboratory. After acclimation to experimental conditions, 20 mL of NSW was added to 25 cm^2^ rectangular cell culture flasks (Falcon) with vented caps and subsequently spiked with different concentrations of spherical polystyrene NPs (Spherotech PP-008-10, 50-100 nm) or MPs (Spherotech PP-20-10, 2.29 µm) to produce test concentrations of 0, 0.00005%, 0.0005%. 0.005% and 0.05% w/v in triplicate. Flasks were maintained at 19ºC at a light intensity of 10 µmol photons m^-2^ s^-1^ and placed on an orbital shaker (140 rpm). After 0, 24 and 72 h, a 1 mL sub-sample was collected from each flask and fixed using 0.25% glutaraldehyde and 0.01% pluronic acid as previously recommended.^54^ Fixed samples were analysed by flow cytometry using a Becton Dickinson FACS-Verse cytometer. FITC (488 nm excitation, 530/30 nm emission) and PE (488 nm excitation, 576/26 nm emission) together with FSC and SSC were used to separate the phytoplankton populations. Distinct phytoplankton groups were identified based upon their natural autofluorescence and gated using FACSDiva software. Cell densities were calculated in respect to spiked reference beads at a concentration of 10^4^ beads mL^-1^ (2.2 μm high Intensity fluorescent Nile Red particles, Spherotech FH-2056-2). To identify statistical variations in cell density between untreated control and treated samples for each respective phytoplankton group, two-way T-tests were utilised at each timepoint (p≤0.05).

### 2.4 Phytoplankton culture growth in response to micro- and nano-plastic exposure

To examine the impact of plastic exposure upon the growth of marine phytoplankton, seven species were exposed to NPs (Spherotech FP-00562-2, 50 nm) or MPs (Spherotech FH-2056-2, 2.15 μm) for a period of 72 h. Axenic phytoplankton cultures of all seven species indicated above were routinely grown under optimal growth conditions as described in section 2.2. To establish exposures under simulated natural cell concentrations, cultures were diluted 100X in oligotrophic NSW (Station L4, Plymouth) to a volume of 2 mL held within a 96-well plate and maintained at 23 °C under a constant light intensity of 10 μmol photons m^−2^ s^−1^. NPs or MPs were added to cultures respectively to establish a test concentration of 0.001% w/v.

Phytoplankton were additionally exposed to copper sulphate to achieve a final Cu^2+^ concentration of 0, 0.1, 10 and 10 µM to act as a positive control and examine their response to a known pollutant. Cyanobacteria were additionally treated with lower Cu^2+^ concentrations (0.001-0.05 µM) owing to their increased sensitivity. Treatments were set-up in triplicate and compared to an untreated control grown in the absence of plastics or Cu^2+^. Following 0, 24 and 72 h growth, phytoplankton were monitored by flow cytometry using a Becton Dickinson Fortessa Flow Cytometer. Samples were monitored directly, and phytoplankton gated using their natural autofluorescence using FACSDiva software. The abundance of phytoplankton was calculated relative to reference beads, as described previously. To identify statistical variations in cell density between untreated control and treated cultures for each species, two-way T-tests were utilised at each timepoint (p≤0.05).

### 2.5 Phyto-PAM Photosynthetic Efficiency analysis

The impact of plastic exposure upon the photosynthetic performance of marine phytoplankton was assessed by monitoring photosynthetic efficiency (Fv/Fm) following 72 h incubation. Axenic cultures of *Prochlorococcus, P. tricornutum* and *E. huxleyi* were prepared as described in section 2.2. To establish exposures, cultures were diluted 100X in oligotrophic NSW (Station L4, Plymouth) to a volume of 100 mL held within 250-mL Erlenmeyer flasks. NPs (Spherotech FP-00562-2, 50 nm) or MPs (Spherotech FH-2056-2, 2.15 μm) were added to flasks to establish a test concentration of 0.001% w/v in triplicate. Following exposure, Chlorophyll fluorescence was assessed at the end of the experiments using a PHYTO-PAM^®^ Fluorometer Analyser (Walz, Germany) equipped with an ED-101US/ MP optical unit, without stirring to avoid any bias in fluorescence signal due to movement of cells between dark and illuminated zones in the cuvette.^55^ Fifteen mins before the measurements, samples were dark-acclimated to allow complete oxidation of PSII reaction centres, being F0 the minimum fluorescence yield measured at low intensity of modulated light (<0.3 μmol photons m^−2^ s^−1^); and Fm the maximum fluorescence yield measured when all primary electron acceptors QA of PSII were reduced after a saturation pulse of 2.6 μmol photons m^−2^ s^−1^ during 0.3 s.^56^ The maximal PSII quantum yield (ΦM) of PSII was assessed, reflecting the state of the water photo oxidation process. Background fluorescence of plastics was determined using a blank sample with MPs or NPs added to culture media. An untreated control, where no plastics were added was also monitored. At the end of the experiment, two-way T-tests were carried out to identify statistical variations in photosynthetic efficiency between untreated control and treated cultures for each species (p≤0.05).

### 2.6 Shotgun proteomic analysis of E. huxleyi exposed to nano- and micro-plastics

As a result of earlier works (section 2.4), the coccolithophore *E. huxleyi* was identified to be particularly sensitive towards plastics exposure. Shotgun proteomic analysis was utilised to identify molecular features of NPs (Spherotech FP-00562-2, 50 nm) or MPs (Spherotech FH-2056-2, 2.15 μm) exposure upon the cellular proteome of *E. huxleyi*. Triplicate untreated and treated *E. huxleyi* cultures were obtained from photosynthetic efficiency experiments described above (section 2.5). Briefly, axenic *E. huxleyi* culture was diluted 100X in oligotrophic NSW (Station L4, Plymouth) to a volume of 100 mL as described above and exposed to NPs or MPs at a concentration of 0.001% w/v, together with non-treated controls, all in triplicate. Following 72 h exposure, samples were immediately placed on ice to halt cellular activity. Subsequently, 100ml samples were centrifuged (4000 xg) for 10 mins at 4μC to form a cell pellet. Cell pellets were immediately flash-frozen on dry ice and stored at -20μC. After allowing cell pellets to thaw at room temperature, samples were resuspended in 1X LDS buffer (ThermoFisher) containing 1% beta-marcaptoethanol. To aid cell lysis, cell pellets were subjected to three cycles of 5 mins sonication (Branson 1210 Sonicator, 40 kHz), followed by incubation at 95μC and a short vortex. The resuspended cell pellets were then run on a precast NuPage 4-12% Bis-Tris polyacrylamide gel at 200V for a period of 5 min.^57, 58^ Gels were subsequently stained using SimplyBlue SafeStain (Invitrogen), and bands containing the cellular proteome were isolated and stored at -20 μC.^58^ In-gel trypsin digestion and peptide recovery^59^ was carried out to prepare samples for nanoLC-ESI-MS/MS analysis. An UltiMate 3000 RSLCnano with Orbitrap fusion (Thermo Scientific) equipped with a 120 min LC separation on a 25 cm column was used to obtain RAW mass spectral files, as previously described.^60^ Subsequently, MaxQuant version 1.5.5.1^61^ was used for peptide identification and protein label-free quantification,^62^ using default settings and “match between runs” for peptide identification. The Uniprot database for *E. huxleyi* used for peptide identification was downloaded on the 04/02/2017, to which coding domain sequences encoded by mitochondria and chloroplasts were included. Downstream analysis was carried out using Perseus version 1.5.5.3, after filtering of the data to remove potential contaminants.^58, 63, 64^ Two-way T-tests (FDR≤0.05) were utilised to assess significant alterations in protein expression between treated and control samples (volcano two-way T-test, p≤0.05). Relative abundance of individual proteins was normalised to peptide length^58^ and utilised to calculate relative abundance of functional protein groups: basic cellular processes; carbon fixation; central metabolism; energy production and conversion; inorganic nutrient processing; other; oxidative stress; photosynthesis; membrane transport; and uncharacterised. Two-way T-tests (p≤0.05) were utilised to identify significant alterations in relative abundance of functional groups between treated and untreated cultures.

### 2.7 Imaging of plastic-cell interactions using confocal microscopy

The occurrence of plastic-cell hetero-aggregation was examined by carrying out fluorescent microscopy upon the coccolithophore *E. huxleyi* exposed to spherical polystyrene NPs (Spherotech PP-008-10, 50-100 nm) or MPs (Spherotech PP-20-10, 2.29 µm) following 72 h exposure. In this work cell-dense cultures (10^6^ cells mL^-1^) were utilised to facilitate imaging, exposed to NPs at a concentration of 0.05% w/v. 2 mL cultures were added to a 96-well plate and maintained under at 23 °C under a constant light intensity of 10 μmol photons m^−2^ s^−1^. After 72 h, 20 µL sub-samples were collected from the bottom of culture flasks and transferred to a glass microscope slide which was immediately imaged using a Leica TCS SPE confocal microscope (Leica Microsystems). Images were obtained using *E. huxleyi* chlorophyll autofluorescence (Ex/Em 440/640-690 nm) and processed using the Leica application suite software (Leica Microsystems). In addition, sub-samples were collected and fixed for flow cytometric analysis, as described above (section 2.2) to monitor changes in the planktonic free-living population.

## 3. Results and Discussion

### 3.1 Characterising the natural community response to micro- and nano-plastic exposure

Exposure of natural marine communities to NPs and MPs revealed significant impacts upon phytoplankton growth related to cell size, being larger phytoplankton more susceptible to plastic exposure than smaller phototrophs. This result is unexpected given the typical response of microorganisms to toxicants, where smaller organisms possessing relatively large surface-to-volume ratios experience greatest toxicity. However, the relationship between cell volume and extent of cell decline in response to NP or MP exposure was only apparent at supra-environmental values, far exceeding those predicted in the environment. Hence, at environmentally-relevant plastic concentrations, negligible impact of plastic exposure upon natural marine photosynthetic communities was observed.

To date, the potential impacts of small ubiquitous plastic debris (<1 mm) upon marine phytoplankton remained uncertain.^29, 30^ To address this, we exposed a natural marine community to NPs (∼50 nm) and MPs (2.29 µm), representing those small plastic particles believed abundant in the environment.^6, 20^ NSW was incubated with NPs or MPs at a range of concentrations (*i*.*e*., 0.00005-0.05% w/v, equating to 0.5-500 mg L^-1^) and phytoplankton growth was monitored by flow cytometry for a period of 72 h (Fig 1). The abundance of distinct groups (*i*.*e*., picocyanobacteria and eukaryotes) was recorded based on their natural autofluorescence and cell morphology (Fig SI.1).

**Figure 1.**
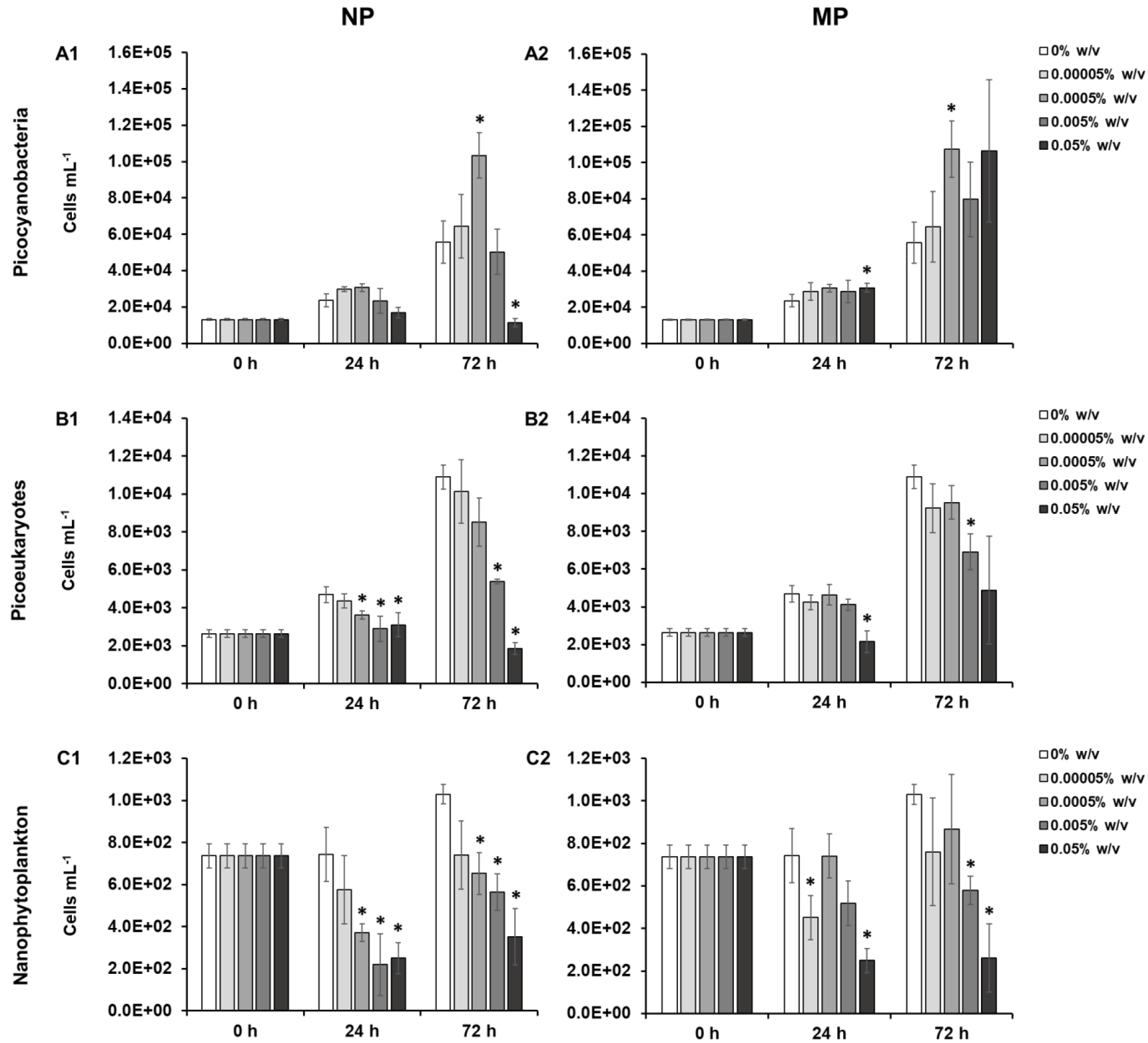
Flow cytometric monitoring of natural phytoplankton communities exposed to 1) nanoplastics (NPs), or 2) microplastics (MPs) at a concentration of 0.00005-0.05 w/v for a period of 72 h. The impact of exposure on abundance of three distinct phytoplankton groups; picocyanobacteria, picoeukaryotes and nanophytoplankton is displayed in panels A-C. Data is presented as the mean ± standard deviation of triplicate samples. Markers indicate where two-way T-tests identified cell density to significantly vary between the untreated control and treated cultures at each timepoint (p≤0.05), respectively.

Based on their natural autofluorescence, three distinct phototrophic groups were identified and monitored by flow cytometry: picocyanobacteria identified mainly as *Synechococcus sp*. (Cyano), picoeukaryotes (Pico-Euk) and nanophytoplankton (Nano-Phyt). At 0 h Cyano were most abundant (1.3×10^4^ cells mL^-1^), followed by Pico-Euk (2.6×10^3^ cells mL^-^1) and Nano-Phyt (7.4×10^2^ cells mL^-1^), respectively.

In NP exposures, no impact upon Cyano growth was recorded after 24 h (Fig 1 A1). However, following 72 h in response to 0.05% w/v NPs, Cyano cell density was reduced approximately 79% compared to the untreated control (two-way T-test, p≤0.05). Previous evidence of cyanobacterial decline in response to NP exposure has been recorded.^39, 40^ When incubated with 100 nm PS NPs (0.0005% w/v, 5 mg L^-1^) cyanobacteria were recorded to form aggregations with particles and settle out of the water column, representing a reduction in the planktonic population; however no adverse impact upon cell viability was recorded as cyanobacteria remained alive during this process.^39^ It is likely that a similar process is responsible for the significant decline in the Cyano population at high NP concentrations recorded herein. In contrast, by the end of the experiment Cyano in the 0.0005% w/v treatment had grown to cell densities 86% higher than the untreated control (two-way T-test, p≤0.05). As a result of MP exposure, no negative effect upon Cyano growth was recorded at any concentration (Fig 1 A2). Rather, cell density remained on average higher than that of the untreated control throughout the experiment, significant after 24 h in the 0.05% w/v treatment (30% increase relative to the control) and after 72 h in the 0.0005% w/v treatment (93% increase relative to the control) (two-way T-test, p≤0.05).

In response to plastic exposure, Pico-Euk cell density was on average lowered by NPs and MPs at every concentration at both 24 h and 72 h timepoints (Fig 1B). This decline was more severe in NP treatments. Here, a significant decrease in Pico-Euk cell density was recorded after 24 h in NP treatments ≥0.0005% w/v, resulting in a 23-60% decrease compared to the untreated control (two-way T-test, p≤0.05). This decline continued throughout the experiment, causing a 51% and 83% decline in the 0.005% and 0.05% w/v NPs treatments respectively after 72 h (two-way T-test, p≤0.05). In response to MPs, significant declines in Pico-Euk were recorded after 24 h in the 0.005% w/v treatment (37%) and 0.05% w/v treatment after 72 h (54%) (two-way T-test, p≤0.05). In-line with the decline in smaller eukaryotic members, Nano-Phyt also experienced significant decreases in growth in response to both NPs and MPs (Fig 1C). Similarly, concentrations ≥0.0005% w/v NPs drove significant declines in large-sized Nano-Phyt populations after 24 h, resulting in decreases of 50-70% compared to the untreated control (two-way T-test, p≤0.05). At this timepoint, MPs at the 0.00005% and 0.05% w/v concentrations also drove relative cell declines of 39% and 67% respectively (two-way T-test, p≤0.05). The significant declines observed in NP treatments continued to the end of the experiment, reaching a maximum decrease in cell density of 66% compared to the untreated control (two-way T-test, p≤0.05). At this final timepoint, MP exposure >0.005% w/v also drove significant declines of up to 75% (two-way T-test, p≤0.05). Negative impacts of plastic exposure have previously been recorded in both algal and diatom species, discussed in greater detail in the following section.^34, 36-38, 65, 66^

It appears from natural community exposure that smaller microorganisms (*i*.*e*., Cyano) are less affected by plastic exposure than larger eukaryotic members of the phototrophic community. Significant declines in the Cyano population are only recorded at extremely high concentrations where reductions in cell density likely arise from co-aggregation with plastics and subsequent precipitation out of the water column.^43, 67^ Such effects have previously been recorded in cyanobacterial exposure to metal oxide nanomaterials.^68, 69^ Interestingly, evidence that presence of plastics can enhance the growth of picocyanobacteria was recorded in a number of treatments regardless of plastic size. Given that picocyanobacterial members dominated the phototrophic community, total cell density appears little affected by plastic exposure. However, this effect masks the negative effects experienced by larger eukaryotic members, which in terms of biomass can represent the equivalent of up to thousands of cyanobacteria. Indeed, larger eukaryotic phototrophs appear more susceptible to plastics, experiencing significantly reduced growth in response to both NP and MP treatments.

### 3.2 Investigating the impact of micro- and nano-plastics upon growth of marine phytoplankton

The enhanced susceptibility of larger phytoplankton to small plastic particles revealed during community exposure was subsequently confirmed by laboratory exposure of axenic phytoplankton cultures towards NPs and MPs. A total of seven phytoplankton species were selected for study, thus widening our knowledge on the possible effects of these contaminants, particularly in terms of NPs for which data is limited. Model species represented four taxonomic groups: cyanobacteria, green algae, diatoms and coccolithophores. In addition, to act as a positive control phytoplankton were exposed to dissolved Cu^2+^, a well-known marine contaminant.^70, 71^ Cultures were diluted in oligotrophic NSW to achieve natural-relevant cell concentrations to best represent natural conditions and exposed to plastics (0.001% w/v) or Cu^2+^ (0-10 µM), after which populations were monitored by flow cytometry for a period of 72 h (Fig SI.2).

Interestingly, in-line with results obtained during community exposure (section 3.1), experimentation revealed both positive and negative impacts of plastic exposure upon phytoplankton growth (Table SI.1). Again, we observed that larger cells appeared to suffer greater adverse effects than their smaller counterparts when exposed to plastic particles as shown by the higher percentage change in cell density relative to the untreated control when plotted against cell volume following NP (Fig 2A) or MP (Fig 2B) exposure. Cu^2+^ was included as a control for the expected cell size-concentration effect of toxicity whereby smaller cells display highest sensitivity towards exposure, owing to their relatively high surface-to-volume ratios.^72, 73^ As expected, cell with the lowest cell volumes exhibited highest sensitivity to Cu^2+^ (Fig 2C). Here, EC_50_ values ranged from 0.05 µM in the cyanobacteria *Prochlorococcus* and *Synechococcus*, to 6.65 µM in the diatom *T. pseudonana* (Table SI.1).

**Figure 2.**
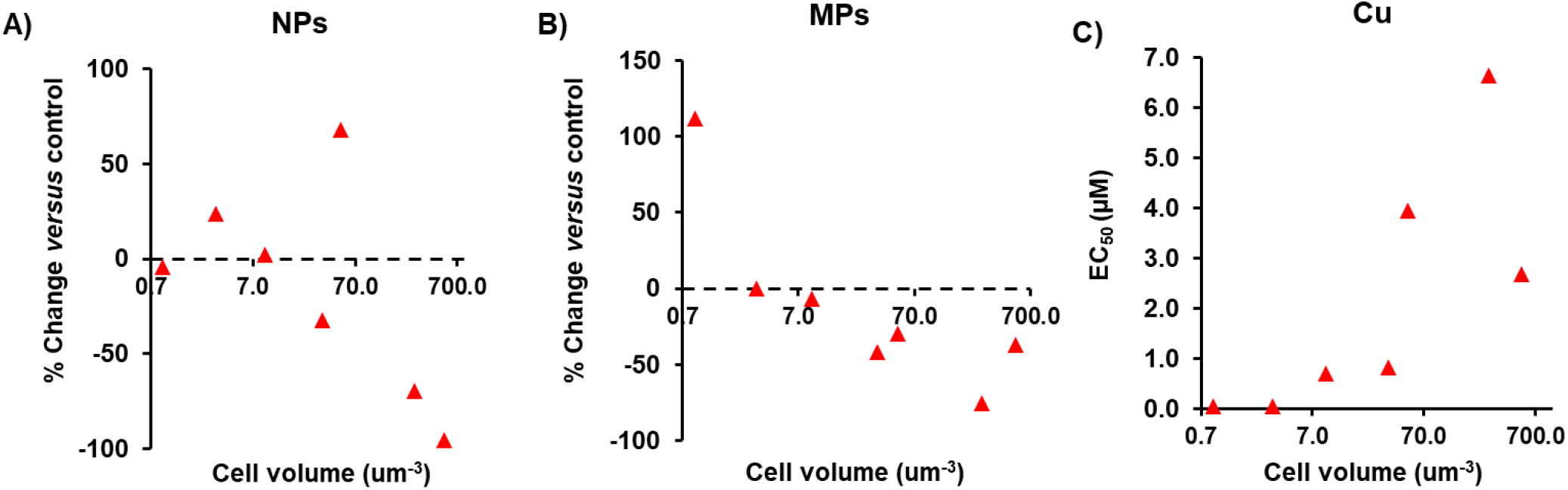
Relationship between cell volume and associated impact of A: NPs (0.001% w/v); B: MPs (0.001% w/v); and C: Cu^2+^ (0-10 µM) exposure upon phytoplankton growth. Seven phytoplankton species were exposed to plastics: *Prochlorococcus sp*. MED4 (0.9 µm^3^), *Synechococcus sp*. WH7803 (3.1 µm^3^), *Ostreococcus tauri* OTH95 (9.2 µm^3^), *Micromonas sp*. CMP2709 (33.5 µm^3^), *Phaeodactylum tricornutum* CCMP2561 (50.0 µm^3^), *Thalassiosira pseudonana* CCMP1335 (268.1 µm^3^) and *Emiliania huxleyi* CCMP 1516 (523.6 µm^3^).

Our findings suggest that the response of phytoplankton to plastic particles is largely taxa-specific, and appears related to cell size, but may also arise as a consequence of varied morphology and physiology. To assess any significant impact of plastic exposure upon phytoplankton growth, two-way T-tests were carried out to compare cell densities of untreated control cultures and those exposed to plastics (0.001% w/v) after 72 h (Fig SI.2). Notably, the coccolithophore *E. huxleyi* (Fig SI.2G), representing the largest organism tested (∼524 µm^3^), appeared particularly susceptible to plastic exposure. This species was significantly reduced by NPs, resulting in 95% loss of the population compared to the untreated control (two-way T-test, p≤0.05; Fig 2A). Although not statistically significant, MP exposure in this species also drove a decline in cell density of approximately 37% compared to the untreated control (two-way T-test, p=0.16). Little work has been carried out examining the impact of plastic exposure towards coccolithophores, however, our results suggest this group may be particularly sensitive to plastic exposure at high concentrations. These species play a key role in the marine carbon pump, both through their photosynthetic and calcification activities,^74, 75^ hence their enhanced susceptibility towards plastic is a concern and requires greater investigation at environmentally-relevant plastic concentrations.

In previous work, diatoms have been identified as being sensitive to NP exposure.^31-35^ For example, following 96 h exposure to 50 nm PS-NH_2_ NPs (0.0005% w/v, 5 mg L^-1^) *Chaetoceros neogracile* was observed to experience a 62% decline in growth when incubated during the exponential growth phase.^36^ Diatom species play a significant role in marine ecological functioning, contributing ∼20% towards global primary productivity.^76^ In our work, the diatom *T. pseudonana*, representing the second largest species tested here (∼268 µm^3^), also suffered declines in average cell density in response to both MPs and NPs, resulting in a respective decline of 75% and 70% in cell density (Fig 2A-B and Fig SI.2F). However, due to variation between replicates this was not statistically significant (two-way T-test, p=0.16; 0.18). Unexpectedly, significantly enhanced growth following plastic exposure was also observed in the *P. tricornutum* (∼50 µm^3^), appearing to be an outlier of the general trends observed (Fig 2 and Fig SI.2E). Here, cell density of NP-treated cultures increased significantly by 68% relative to the control (two-way T-test, p≤0.05). Evidence of improved phytoplankton growth in presence of plastics has previously been recorded in the freshwater microalga *Raphidocelis subcapitata*.^77^ Hence, it appears that despite cell size playing an influential role in determining the likely impact of small plastic particles upon phytoplankton, there is evidence of species-specificity in response. As a result, differential sensitivity displayed by varying species may cause alterations to community structure and functioning, however this is only expected in high levels of plastic pollution given the high concentrations examined herein. Despite the enhanced growth of *P. tricornutum* recorded, evidence suggests that diatoms may be more susceptible to small plastic particles than other phytoplankton groups.^34^ This is believed to be attributed to morphology^34, 37^ and the ability of diatoms to produce sticky exopolymeric substances (EPS), increasing the likelihood of adherence of NPs to cell structures.^34, 43^ However, interestingly Grassi *et al*. (2020) found no adverse effect of negatively-charged PS-COOH NPs (0-0.01% w/v, 0-100 mg L^-1^) upon *P. tricornutum* after 72 h exposure in cultures supplemented with EPS, characterised by a reduction in intracellular ROS generation.^78^ The presence of EPS was found to reduce aggregation of NPs, which were recorded to aggregate extensively upon entry into saline media, likely due to adsorption of EPS biomolecules onto the NP surface which is also hypothesised to enhance scavenging of harmful ROS.^78^ Therefore, EPS may also mitigate plastic toxicity as well as facilitate physical cell-plastic interactions.

Green algae appear susceptible to both NP and MP exposure, however, again this is largely taxa specific as recorded in our work presented herein and in the literature.^34^ The green algae, *Ostreococcus* suffered no negative effect of plastic exposure during experimentation (Fig SI.2C), whereas average cell density of *Micromonas* was reduced by 32% and 42% in the presence of NPs and MPs respectively by the end of the experiment (Fig SI.2D). This result can be expected given that cell volume of *Micromonas* is approximately three-times that of *Ostreococcus* which represents the smallest known eukaryotic phototroph.^79^ In previous research similar alterations in plastic impacts upon algal taxa are observed, with smallest cells also experiencing lowered sensitivity to exposure.^34^

NP exposure exerted no negative impact upon the smallest phytoplankton species tested: *Prochlorococcus* (Fig SI.2A) and *Synechococcus* 7803 (Fig SI.2B). In fact, cell density of *Prochlorococcus* was observed to display enhanced growth in the presence of MPs, resulting in a significant 112% increase in cell density (two-way T-test, p≤0.05).

### 3.3 Mechanistic understanding of the adverse effect of MP and NP on large-size phytoplankton

#### a) Examining the impact of plastic exposure upon the photosynthetic efficiency of marine phytoplankton

Primary productivity in the marine environment contributes approximately 50% of global oxygen production,^28^ hence improving our understanding of the likely impact of plastic exposure upon the photosynthetic performance of marine phytoplankton is key. In previous research, disruption of normal photosynthetic function has been highlighted as a feature of exposure towards plastic particles.^29, 36, 38, 80^ To investigate any alterations to photosynthetic function, a PHYTO-PAM fluorimeter was utilised to assess the photosynthetic efficiency (Fv/Fm) of four phytoplankton species grown in presence of NPs or MPs (0.001% w/v) within oligotrophic NSW (Fig 3). Exposure to both sizes of plastic resulted in a decline in average photosynthetic efficiency in all three species tested. In two species, *P. tricornutum* and *E. huxleyi*, only NP exposure caused a significant decline in photosynthetic efficiency compared to the untreated control (two-way T-test, p≤0.05). Rather, in *Prochlorococcus* MP exposure significantly reduced photosynthetic performance (two-way T-test, p≤0.05). For *Prochlorococcus* and *P. tricornutum*, average declines in Fv/Fm ranged 0.07-0.13, representing declines of 15-35% in photosynthetic efficiency compared to the control. However, adverse effects on photosynthetic efficiency appeared most severe for *E. huxleyi*, shown above to be particularly susceptible to NP exposure (section 3.2). Here, Fv/Fm was recorded to decline from 0.39 in untreated cultures to 0.20 and 0.06 upon the addition of MPs and NPs, representing respective declines of 48% and 84% compared to the control.

**Figure 3.**
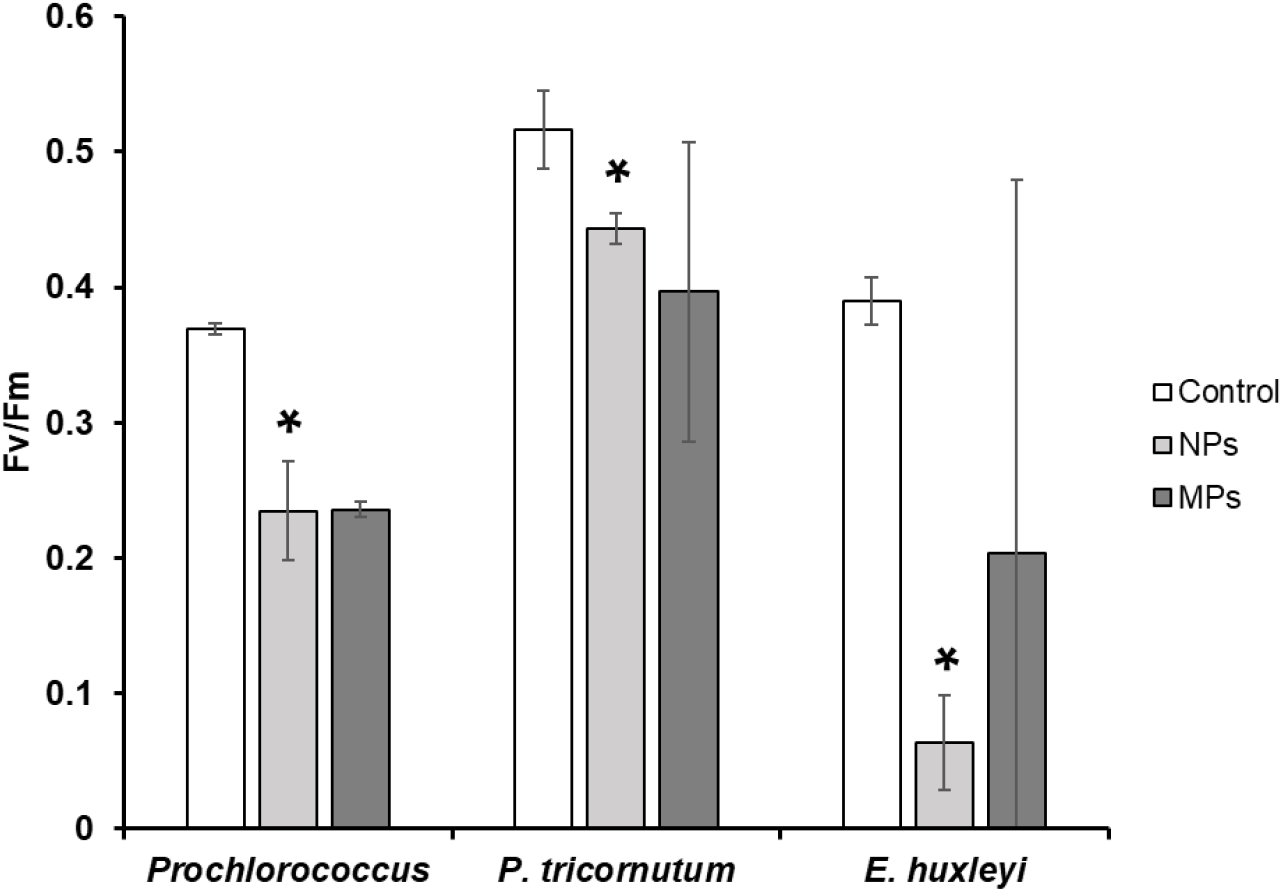
Photosynthetic efficiency (Fv/Fm) of marine phytoplankton exposed to NPs or MPs (0.001% w/v) as measured using a Phyto-PAM. Data is presented as the mean ± standard deviation (n=3). Markers indicate where photosynthetic efficiency significantly differed between control and treated cultures (two-way T-test, p≤0.05).

A reduction in photosynthetic efficiency has previously been recorded in studies examining the impact of plastic exposure upon phytoplankton growth.^29, 31, 81, 82^ González-Fernández *et al*. (2020), reported that the photosynthetic machinery of the diatom *C. neogracile* was significantly altered after exposure to NPs.^80^ In this work, a range photosynthetic pigments were significantly reduced,^80^ believed to be a rapid photoprotective response to stress.^80, 83^ Plastic exposure is often associated with a reduction in chlorophyll content.^31, 32, 36, 38, 66^ For example, in response to 50 nm PS-NH_2_ both chlorophyll content and photosynthetic efficiency were reduced 33% and 13% respectively in the *C. neogracile*, resulting in a growth reduction of up to 62%.^36^ Similar results have also been obtained during work examining freshwater species.^84^ It is believed that such damages to photosynthetic pigments may arise from an increase in intracellular ROS which may damage pigment structure, or as a result of hetero-aggregation the entry of light and nutrients into the cell is compromised, thus reducing energy available for pigment synthesis or simply the need of sustaining such extensive photosynthetic machinary.^29, 43, 85^ Bhattacharya *et al*. 2010, utilised a CO_2_ depletion assay to demonstrate the ability of hetero-aggregations of plastic particles to hinder algal photosynthesis. Despite the evidence of damages to photosynthetic function of marine phytoplankton available, results vary, with evidence of enhanced photosynthetic performance also recorded.^43, 86^ In response to polyethylene MPs, *D. salina* displayed improved growth associated with an increase in chlorophyll content and improved photosynthetic efficiency, believed due to the release of trace levels of additives added to the plastic particles.^86^ Related to this, photosynthetic activity has also been recorded to improve when phytoplankton were exposed to expanded polystyrene leachate in four taxa.^87^ Further investigation is required to fully understand the likely impact of plastic exposure upon photosynthetic processes, particularly under environmental conditions.

#### b) Shotgun proteomic analysis of E. huxleyi in response to micro- or nano-plastic exposure

In response to plastic exposure, microorganisms may alter the regulation of specific cellular processes in order to adapt to- or mitigate stress. Plastic exposure has previously been associated with DNA damage^38^ and altered gene function^88, 89^ in marine phytoplankton. Owing to the apparent enhanced susceptibility of *E. huxleyi* towards plastics exposure observed during flow cytometric monitoring (section 3.2) and examination of photosynthetic performance (section 3.3a), this species was selected for further investigation of potential toxic mechanisms *via* shotgun proteomic analysis.

Following quality filtration of data, a total of 188 proteins in the cellular proteome of *E. huxleyi* were used for downstream analysis. Statistical analysis revealed nine individual proteins to be significantly altered in the presence of plastic particles (volcano two-way T-test, p≤0.05). All significant proteins were identified in the NP treatment (Fig SI.3), each being significantly less abundant than in control cultures. Interestingly, no significant proteins were identified in the MP treatment, in accordance with earlier experiments displayed above which also reported no significant impact of MPs on growth or photosynthetic efficiency of *E. huxleyi*. Such findings are in accordance with previous research which observe enhanced impacts of plastic particles upon phytoplankton with decreasing particle size.^44, 86^ Of the nine significant proteins identified, seven were annotated with a known function (Fig 4): (A) ATP synthase subunit alpha, (B) ATP synthase gamma, (C) Light harvesting protein, (D) Photosystem I P700 chlorophyll a apoprotein A1, (E) Ribulose bisphosphate carboxylase large chain (RuBisCo), (F) Transketolase and (G) the putative EF-1 alpha/Tu like protein; all representing key proteins in the three relevant biological functions: energy production, photosynthesis and carbon fixation. The Photosystem I P700 chlorophyll *a* apoprotein A1 involved in electron transport was decreased -2.48 Log2FC, whilst the light harvesting protein, usually part of a photosystem to enhance light capture was reduced by -3.47 2LogFC. Hence, the ability of *E. huxleyi* to effectively capture and utilise light energy may be compromised by the presence of NPs. This finding is in accordance with results obtained with the Phyto-PAM, displaying a significant decrease in photosynthetic efficiency in the presence of NPs. Reduced expression of chloroplastic genes was previously recorded in algal exposure to plastics in freshwater species.^89^ Notably, the key RuBisCo protein was downregulated approximately -2.74 Log2FC, which alongside alterations in Transketolase abundance is likely to reduce carbon fixation, which in this ecologically important species may have environmental implications.^30^ In addition, through the decreased abundance of ATP synthase alpha and gamma, ATP production may be reduced.

**Figure 4.**
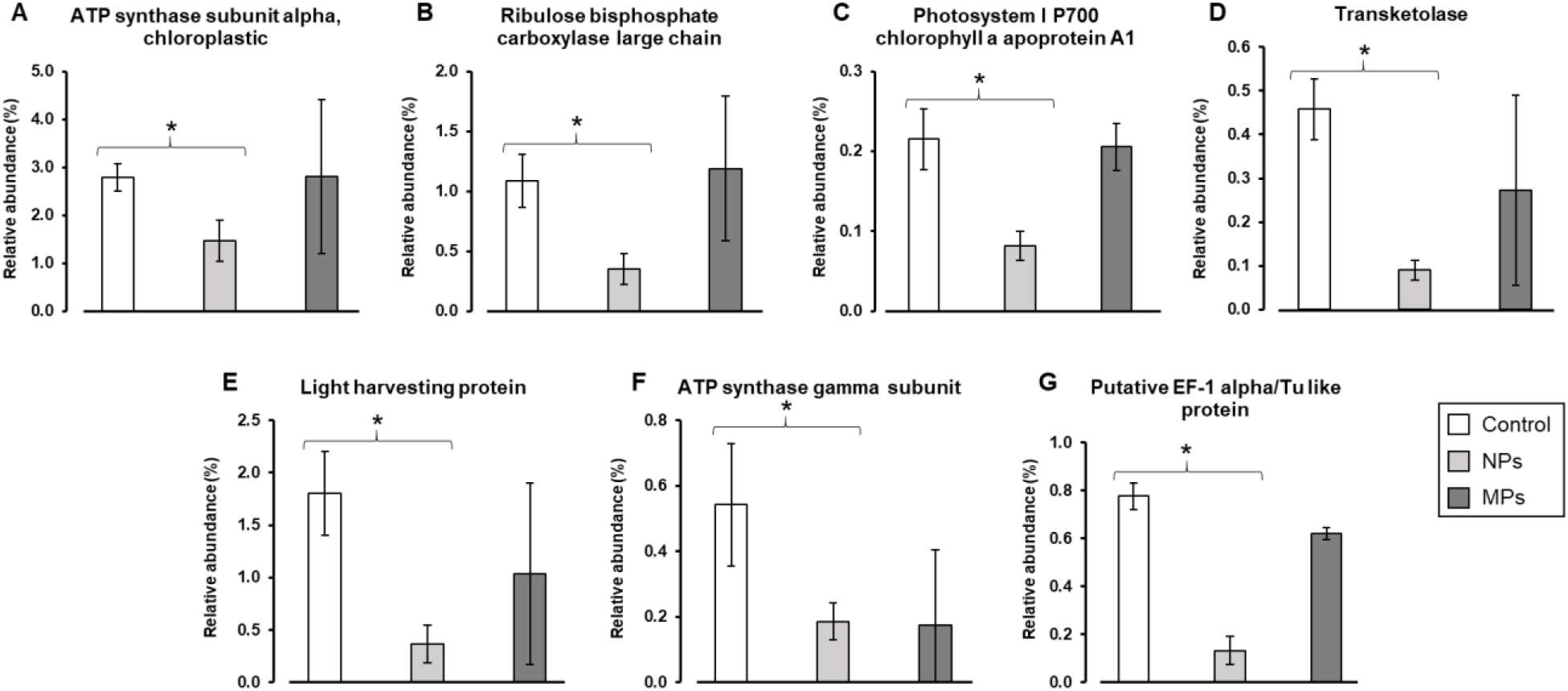
Relative abundance of significant proteins identified during shotgun proteomic analysis of *E. huxleyi* exposed to NPs at a concentration of 0.001% w/v. Data is presented as the mean ± standard deviation.

Trends observed at the individual protein level are strengthened upon examining the relative abundance of functional proteins groups in each treatment (Fig SI.4). Once more, significant alterations are only observed in the NP treatment, where those proteins associated to photosynthesis and energy production are significantly decreased ∼14-10% and ∼9-4%, respectively (two-way T-test, p≤0.05). Also of note, proteins involved in central metabolism were on average more abundant in plastics treatments, ∼19% and ∼18% in the presence of NPs and MPs respectively, compared to ∼12% in the untreated control. Alterations to metabolic function have previously been recorded in cyanobacteria and diatom species in response to NPs.^36, 40, 90^

Oxidative stress has previously been recorded as a feature of plastics exposure towards marine microbial organisms.^29, 36-38, 81, 91^ In our work, oxidative stress proteins were on average more abundant in the NP treatment compared to the control (0.23% *versus* 0.08%), characterised primarily by an increase in superoxide dismutase. The induction of superoxide dismutase and malondialdehyde is described as a feature of plastic exposure in marine phytoplankton, a response that appears less severe in freshwater species.^29^ The increase in such antioxidant proteins is believed to arise due to the presence of ROS, or as a consequence of membrane damage caused by physical contact with plastic particles.^92^ Oxidative stress arising from ROS generation appears a feature of NP exposure which may be enhanced at small particle sizes due to increased surface-area to volume and likelihood of internalisation. For example, after just 0.5 h exposure to PS-NH_2_ (50 nm) and non-modified PS (55 nm) NPs (0.008% w/v, 80 mg L^-1^) the intracellular ROS content of the marine proteobacterium *H. alkaliphile* was recorded to increase 4.4- and 7.7-fold respectively; whereas, no significant impact was observed in response to micron-sized PS.^91^ Significant increases in intracellular ROS have also been recorded in several diatom species.^36-38^ *P. tricornutum*, displayed significant increases in intracellular ROS in response to PS NPs (50 and 100 nm) during 72 h exposure where negative impacts upon growth were also recorded.^38^ Similarly, *C. neogracile* experienced increased intracellular ROS generation up to 48% in response to PS-NH_2_ (50 nm), reducing over time to 22% after 72 h.^36^ Indeed it appears that intracellular ROS generation following NP exposure occurs most rapidly during early stages, after which ROS content is reduced over time.^36, 38, 91^ As well as increases in intracellular ROS content, NP exposure has been observed to enhance ROS generation outside of cells. Bellingeri *et al*. (2020) recorded significant increase in extracellular ROS content in *S. marinoi* cultures exposed to PS-COOH NPs (90 nm) at a concentration of 0.005% w/v (50 mg L^-1^). ROS production likely to cause lipid peroxidation, membrane instability, as well as damage to photosynthetic machinery and processes,^36^ perhaps playing a role in phototoxicity previously described. Interestingly, the presence of supplementary EPS derived from diatom species has been observed to significantly reduce ROS production in the presence of PS-COOH NPs (60 nm) at similar concentrations (0.001% and 0.005% w/v, 10 and 50 mg L^-1^).^78^ This suggests that EPS displays antioxidant activity,^78^ or by its presence mitigates the production of ROS by plastics. In the natural environment where concentration of EPS and related material is likely higher, NP-mediated ROS generation may be mitigated.

#### c) Hetero-aggregation of plastics and phytoplankton

Confocal microscopy confirmed the occurrence of hetero-aggregation between large phytoplankton such as *E. huxleyi* and plastic particles during laboratory exposure. Physical interaction between phytoplankton cells and plastic particles has been regularly reported.^33, 34, 36-39, 88, 93, 94^ Typically, advanced imaging techniques (*i*.*e*., transmission/scanning electron microscopy, fluorescent microscopy) have typically been used to assess this phenomenon,^33,37^ with flow cytometric analysis also identified as a tool for this purpose.^38, 43^ Hetero-aggregation is commonly associated with a decline in the phytoplankton population,^33, 94^ as well as damaging cell morphology^37^ and photosynthetic performance.^81^ The direct impact of hetero-aggregation and subsequent co-precipitation and indirect effects upon photosynthetic performance are believed to be responsible for the significant declines recorded in large phytoplankton at supra-environmental plastic concentrations.

Herein, confocal microscopy was utilised to examine the occurrence and likely role of hetero-aggregation in driving significant declines in phytoplankton growth. Cultures of *E. huxleyi*, identified to be particularly susceptible to plastics exposure, were exposed to ∼50 nm NPs and ∼2 µm MPs (0.05% w/v) for a period of 72 h before samples were prepared for imaging. Here, cell dense cultures were used to facilitate the imaging of cells which would be difficult in more dilute cultures. Samples were obtained from the bottom of culture flasks and the occurrence of hetero-aggregation following both NP (Fig 5B) and MP exposure (Fig 5C) was evident. Cells of *E. huxleyi*, identified using the natural autofluorescence of chlorophyll, could be seen entrapped within large aggregates of plastic particles which reached sizes exceeding 50 µm. To confirm that aggregated material did not consist of any other particulate matter, control samples where no plastics were added were also imaged (Fig 5A), displaying no evidence of such material. Alongside imaging, sub-samples of exposed *E. huxleyi* culture were collected for cell enumeration by flow cytometry after 72 h. Here, cell density was recorded to be reduced on average 95% and 43% relative to the untreated control, in response to NPs and MPs respectively (Fig SI.5). As such, it appears feasible to suggest that the cell declines recorded in *E. huxleyi* and other large phytoplankton above in response to plastic particles, are largely driven by hetero-aggregation and this may only occur at the relatively high plastic particle concentrations investigated. Following this, entrapped phytoplankton are removed from the water column via precipitation of aggregated material, as has previously been recorded.^39, 88, 94^ This process, also observed during exposure with metal oxide nanomaterials,^68, 69^ is believed largely responsible for cases of reduced planktonic cell numbers or OD_600_ recorded in toxicity testing.^39, 88, 94^ Hetero-aggregation between plastic particles and phytoplankton may also act to disrupt photosynthetic processes through possible shading effects, also believed a possible feature of nanomaterial exposure during laboratory investigation.^95^ Large aggregates formed upon the cell surface may act to reduce light availability, compromising photosynthetic performance. Such an effect is in-line with results obtained from both photosynthetic efficiency assessment and shotgun proteomic analysis, earlier described (sections 3.3a and 3.3b).

**Figure 5.**
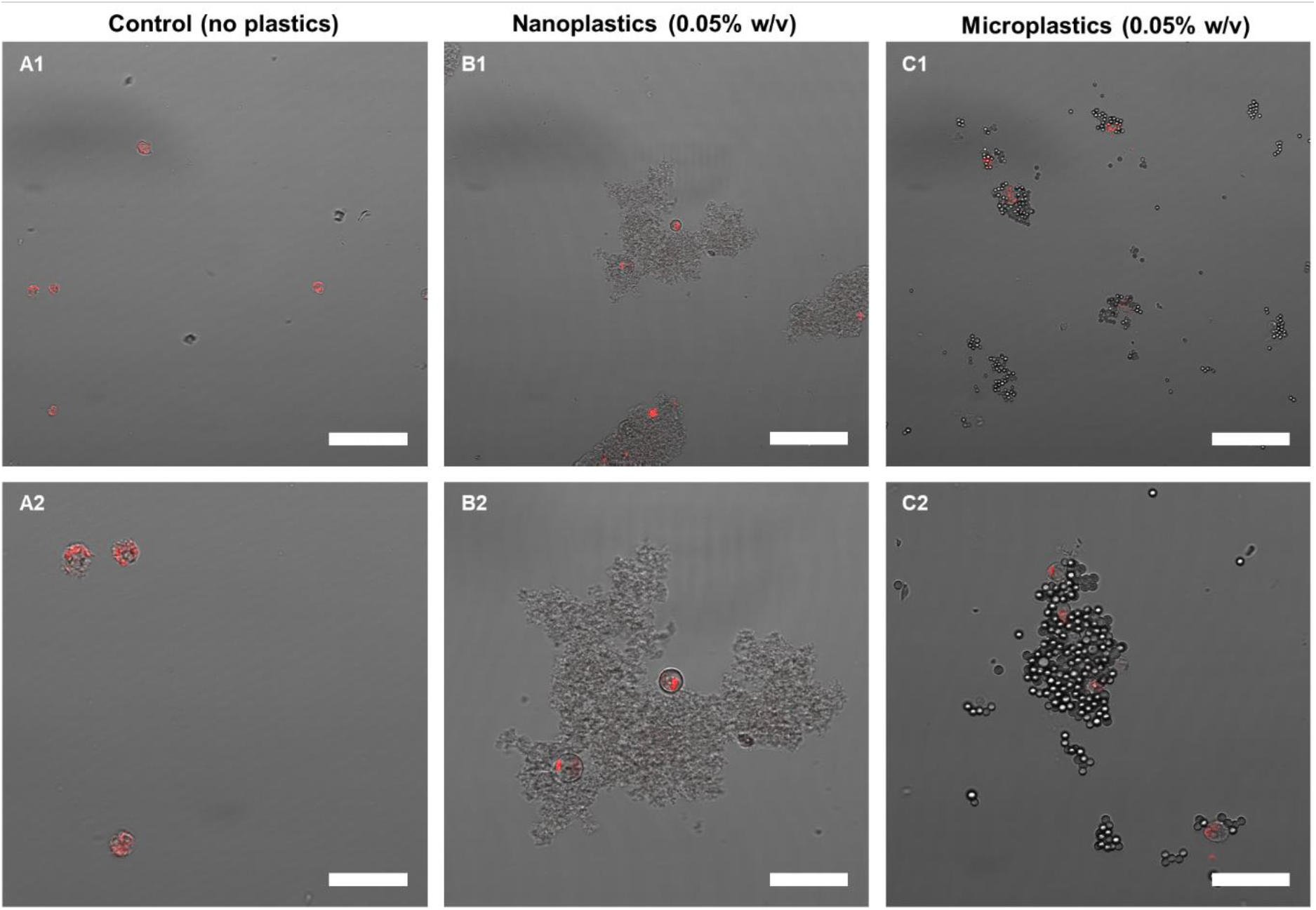
Confocal microscopic imaging of *Emiliania huxleyi* cultures (red) exposed to; A) no plastics; B) nanoplastics; or C) microplastics, added at a concentration of 0.05% w/v for a period of 72 h. Scale bar: 50 µm.

A range of taxa have been adversely impacted by plastics exposure in this manner including dinoflagellates,^88^ cyanobacteria^39^ and heterotrophic bacteria,^94^ in each case following exposure to PS NPs. Such processes may have important ecological consequences.^88, 96^ For example, removal of symbiotic dinoflagellates has been recorded in experimentation with PS NPs (42 nm), which would likely affect their symbiotic relationship with coral species.^88^ Hetero-aggregation between microbial cells and NPs may also facilitate the delivery of co-stressors which are effectively adsorbed and carried by the NP particles.^93^ Likewise, organisms occupying higher trophic levels may ingest such aggregates, facilitating the trophic transfer of plastics through the marine food web.^34^ However, such effects are only likely in areas of extremely high plastic contamination.

According to the manufacturer’s information, the PS particles utilised in our study possess sulfate groups on their surface and are negatively charged. Aggregation of negatively charged NPs upon entry into saline media is widely reported in the literature,^33, 78, 97^ due to maximised particle attachment following the compression of the electrostatic double layer surrounding individual particles in high ionic strength media.^98, 99^ The increased extent of aggregation of negatively charged plastic particles such as those used in our study is likely to have played a key role in the entrapment of *E. huxleyi* within plastic aggregates evident by microscopy. Negatively charged PS-COOH NPs (40 nm) have previously been recorded to aggregate to sizes in the micron range and adhere to marine algal cells.^33^ For positively charged particles such as PS-NH_2_, aggregation with cells may also be facilitated by electrostatic attraction towards negatively charged cell surfaces.^33, 94^

The production of EPS has been recorded as a defensive strategy utilised by microorganisms against nanomaterials,^100^ however these ‘sticky’ structures are thought to increase extent of hetero-aggregation between cells and NPs.^39, 94^ In the case of *E. huxleyi*, coccolith formation and attachment during cell growth is associated with the production of polysaccharides.^101, 102^ The presence of these compounds may enhance the attachment of plastic particles to cells, playing a key role in their decline. Although, as mentioned above presence of EPS can also decrease the extent of aggregation of plastics.^78^ Interestingly, recent findings report an ability of phytoplankton to alter the chemical composition of EPS in the presence of plastic particles, in turn altering their fate and behaviour.^103^ Specific cell structures have been associated with an increased risk from possible physical interaction with plastic particles, which may act to enhance a species’ sensitivity to exposure.^34, 37^ PS NPs were found to freely attach to the fultoportula process of diatoms, which act to form the phytoplankton’s chain-structure.^37^ As a consequence of attachment, NPs were believed to weaken these structures, associated with a reduction in diatom chain length as observed by microscopy.^37^ Further investigation using scanning electron microscopy may provide insight into the physical interaction between plastic particles and the coccoliths of *E. huxleyi*.

It is clear that under laboratory conditions, physical interactions between plastic particles and microorganisms occur freely at the relatively high plastic concentrations that have largely been studied (*i*.*e*., mg L^-1^). It is important that we take such interactions into consideration when evaluating the outcome of exposure and identifying the primary cause of cell decline or reduction in fitness. A consideration of the environmental implications of hetero-aggregation must also take place, as well as examination of the likelihood of plastic particles to interact with other naturally-occurring particulate or organic matter which may act to mitigate cell contact or mitigate toxicity, such as recorded in the presence of EPS.^78^ Of course, due to dilution effects at the lower concentrations predicted in the environment (ng L^-1^ - µg L^-1^)^24^ this process of cell removal is less likely as the rate of encounter between phytoplankton and plastic particles is reduced.

## 4. Conclusions

In this study, we provide new information regarding the impact of nano-(NP) and micro-plastics (MP) upon marine phytoplankton which play key ecological roles. Exposure of both natural marine phytoplankton communities and laboratory cultures revealed that adverse outcomes of exposure to small plastic particles (NP, ∼50 nm and MP, ∼2 µm) are largely related to cell volume and taxonomy. Larger phytoplankton species appear particularly susceptible to plastic exposure, where effects appear enhanced with decreasing particle size. Smaller phytoplankton, on the other hand, appear little affected by exposure to NPs or MPs. Such results do not follow the traditional understanding of toxicity whereby smaller cells display greatest sensitivity due to relatively high surface-area-volume ratios. The coccolithophore *E. huxleyi*, representing the largest organism examined, was found to be most sensitive to plastic exposure, decreasing up to 95% in cell density following 72 h incubation with NPs (50 nm). These findings expand our current knowledge on plastic toxicity, particularly for nanoplastics for which limited data exists, and enables the prediction of the likely impacts of plastic exposure upon various phytoplankton.

Investigation into the drivers of cell declines exerted upon large phytoplankton by plastic exposure revealed hetero-aggregation between cells and large aggregates of plastic particles to be a key mechanism. Cells of *E. huxleyi* could clearly be seen entrapped within aggregates of both NPs and MPs by confocal microscopy. It is believed that the significant cell decline of *E. huxleyi* recorded within laboratory exposure, as well as for other large phytoplankton, largely results from entrapment and subsequent co-precipitation within aggregates of plastic particles, as has previously been demonstrated with metal oxide nanomaterials.^68, 69^ Exposure to NPs was also shown to cause damages to photosynthetic efficiency in *E. huxleyi*, as well as the diatom *P. tricornutum* and cyanobacteria *Prochlorococcus*. Significant impacts upon the cellular proteome of *E. huxleyi* following NP exposure were also recorded, specifically affecting processes of photosynthesis and carbon fixation. Entrapment within plastic aggregates likely reduces light availability due to shading effects and is likely responsible for the adverse effects upon the photosynthetic processes observed.

Global marine microplastic concentrations are predicted in the range of ng-µg L^-1^.^21, 24^ The adverse effects recorded upon phytoplankton presented herein occur only at high concentrations (≥0.0005% w/v, ≥5 mg L^-1^), orders of magnitude above the concentrations of plastic particles believed to be present in the marine environment. The negative effects associated with extensive aggregation of the PS NPs and MPs utilised in our experiments are likely exacerbated at the high concentrations used and closed-system nature of exposures and it is likely that negative effects reported in previous studies, largely carried out at concentrations in the mg L^-1^ range, also result from this process. Hence, in accordance with risk assessments carried out by Everaert *et al*. (2018) the likely risk of small plastic particles towards marine phytoplankton within the natural environment is negligible. Whilst our findings reveal mechanistic insight into the potential effects of plastics upon key phytoplankton species, in the natural environment adverse effects appear limited to heavily polluted zones.

## Acknowledgements

The research was funded through the Newton Fund Latin America Researcher Links Travel Grants (RLTG9-357288064) of the British Council. CJD was supported by the NERC CENTA DTP studentship NE/L002493/1. JAC-O was funded by a NERC Independent Research Fellowship NE/K009044/1, Ramón y Cajal contract RYC-2017-22452 (funded by the Ministry of Science, Innovation and Universities, the National Agency of Research, and the European Social Fund) and project PID2019-109509RB-I00 / AEI / 10.13039/501100011033. PE was funded by Fondecyt Iniciación 11170837, REDI170403 and INACH-RT-12-19. We also thank the technical assistance in imaging by Dr Ramis at the Cellomics Unit (IUNICS, SCT) of the University of the Balearic Islands. In addition, we thank the BBSRC/EPSRC Synthetic Biology Research Centre WISB (grant ref.: BB/M017982/1) for access to equipment.

