## Supplementary Information for "Cell size matters: nano- and micro-plastics preferentially drive declines of large marine phytoplankton due to co-aggregation"

[2] Department of Biology, University of the Balearic Islands, Ctra. Valldemossa, km 7.5. CP: 07122, Palma, Spain.

[3] Instituto de Ciencias Naturales Alexander von Humboldt, Universidad de Antofagasta, Antofagasta, Chile

[4] Instituto Milenio de Oceanografía, Concepción, Chile

### Supplementary Information

#### *SI.1 Flow cytometric analysis of natural phytoplankton communities.*

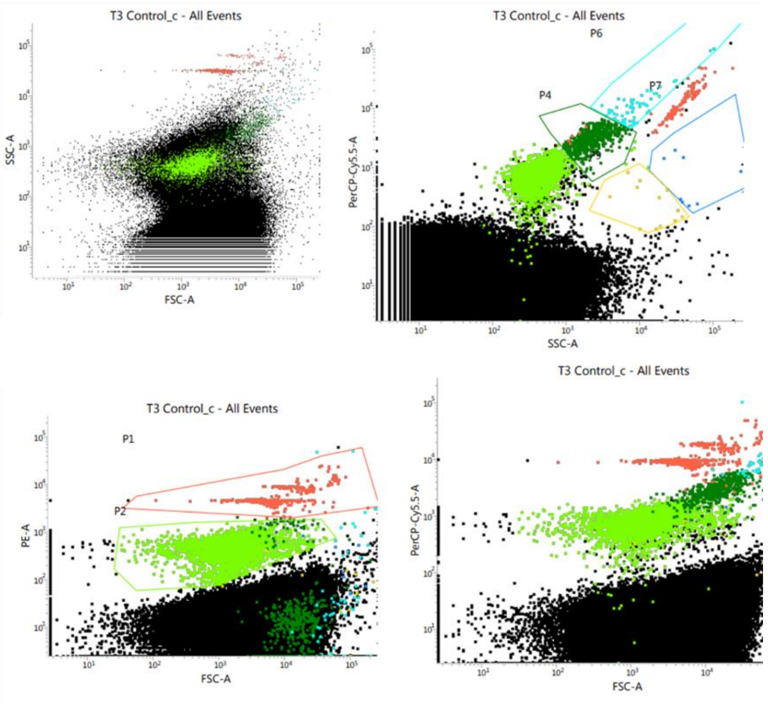


**Figure SI.1.** Gating of phytoplankton groups utilised for determination of respective cell densities in response to nano- and micro- plastics exposure. Light green: Picocyanobacteria; Dark green: Picoeukaryotes; Blue: Nanophytoplankton; Red: reference beads used for calculation of cell density.

### *SI.2 72 h phytoplankton growth*

**Table SI.1.** Summary of data obtained following 72 h exposures of phytoplankton to nano- (NPs) and micro- plastic (MPs) particles, or copper (Cu).

*Significant alteration in cell density between treated and untreated cultures (two-way T-test, p<0.05).

| Species | Cell Volume ($\mu m^3$) | % Change in cell density NPs | % Change in cell density MPs | $EC_{50}$ Cu |
|---|---|---|---|---|
| *Prochlorococcus sp.* MED4 | 0.9 | -4.42% | +112.05%* | >0.05 |
| *Synechococcus sp.* WH7803 | 3.1 | +23.88% | +0.41% | >0.05 |
| *Ostreococcus tauri* OTH95 | 9.2 | +1.98% | -6.43% | 0.70 ± 0.04 |
| *Micromonas sp.* CMP2709 | 33.5 | -32.12% | -41.67% | 0.82 ± 0.57 |
| *Phaeodactylum tricornutum* CCMP2561 | 50.0 | +67.68%* | -29.80% | 3.96 ± 2.82 |
| *Thalassiosira pseudonana* CCMP1335 | 268.1 | -69.68% | -75.19% | 6.65 ± 5.43 |
| *Emiliania huxleyi* CCMP 1516 | 523.6 | -95.06%* | -36.97% | 2.69 ± 0.37 |

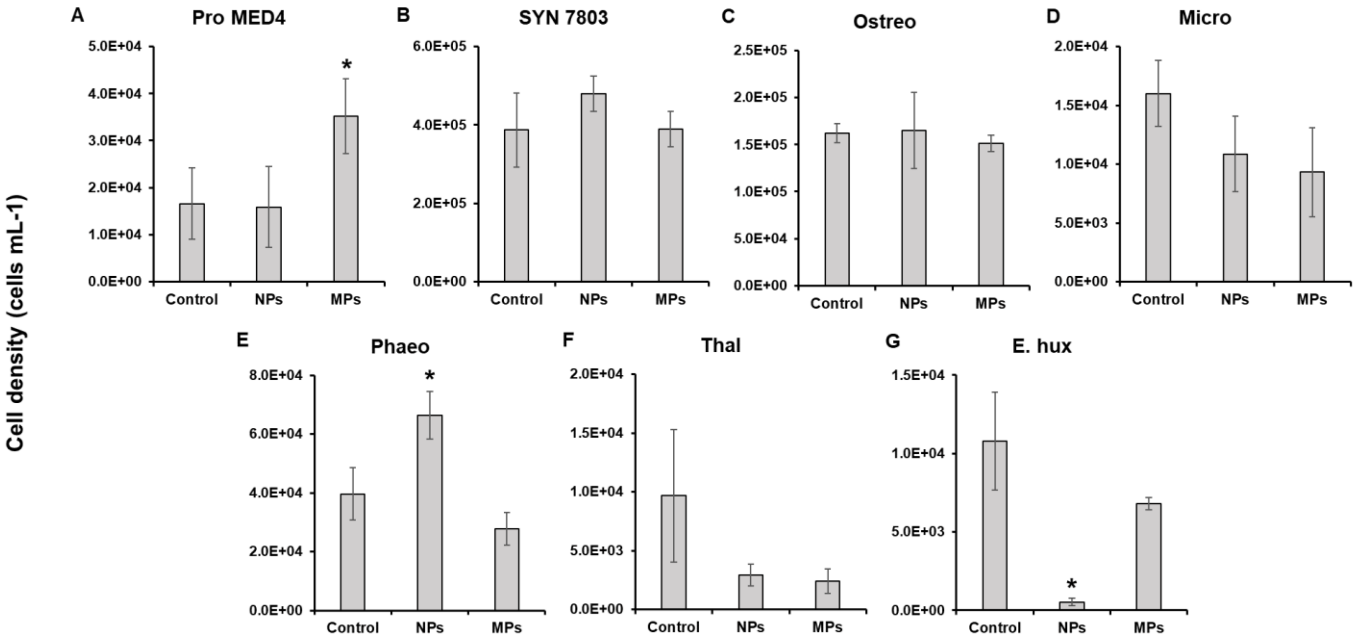


**Figure SI.2.** Cell density of phytoplankton grown in the presence of nanoplastics (NPs) or microplastics (MPs) added at a concentration of 0.001% w/v for a period of 72 h. A: *Prochlorococcus sp.* MED4 (Pro MED4); B: *Synechococcus sp.* 7803 (SYN 7803); C: *Ostreococcus tauri* OTH95 (Ostreo); D: *Micromonas sp.* CMP2709 (Micro); E: *Phaeodactylum tricornutum* CCMP2561 (Phaeo); F: *Thalassiosira pseudonana CCMP1335* (Thal); G: *Emiliania huxleyi* CCMP1516 (E. hux). Data is presented as the mean ± standard deviation of triplicate samples. Stars indicate where two-way T-tests identified cell density to significantly vary between the untreated control and treated cultures at the 72 h timepoint (p≤0.05).

*SI.3 Shotgun proteomic analysis supporting information.*

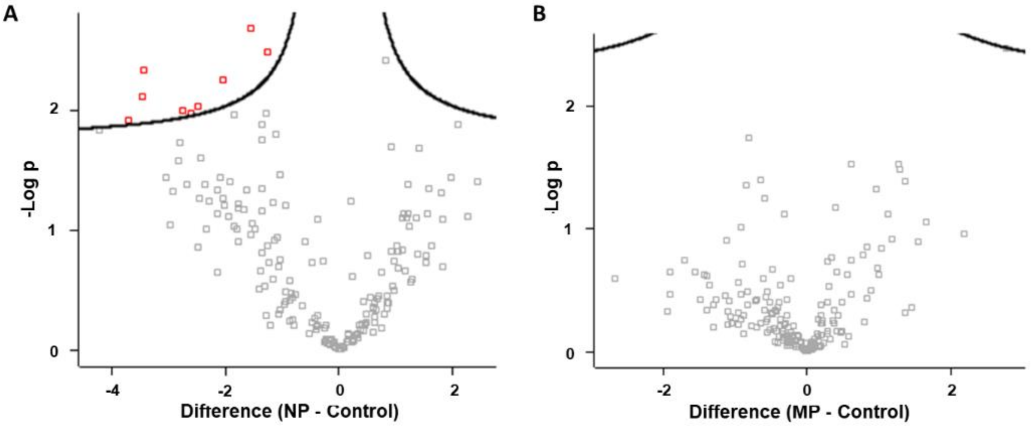


**Figure SI.3.** Volcano plots (T-test; FDR=0.05, S0=0.1) of the cellular proteome of *Emiliania huxleyi* exposed to A) nano- and B) micro- plastics (0.001% w/v). Red markers indicate proteins identified as significantly altering in abundance between control and treated samples (two-way t-test, p≤0.05)

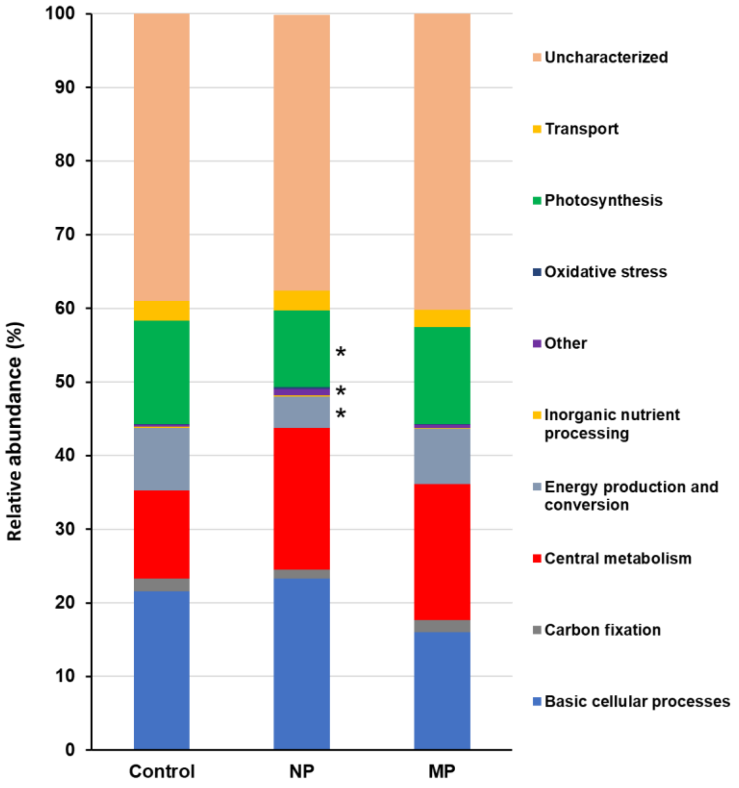


**Figure SI.4.** Relative abundance of protein groups identified in the cellular proteome of *Emiliania huxleyi* exposed to nano- (NP) or micro- plastics (MP) at a concentration of 0.001% w/v. Markers indicate where relative abundance of individual protein groups varies significantly from the untreated control, as identified by two-way T-tests (p≤0.05).

*SI.4 Emiliania huxleyi cell density following 72 h exposure to plastics during imaging experiment.*

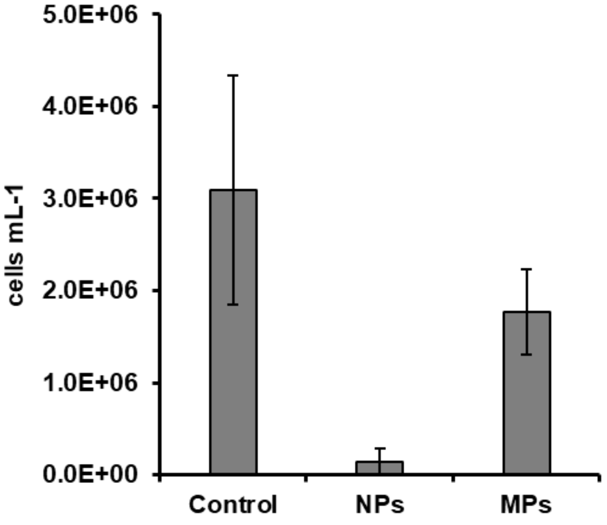


**Figure SI.5.** Cell density of *Emiliania huxleyi* following 72 h exposure to nano- (NPs) or micro-plastics (MPs) at a concentration of 0.001% w/v (n=3). Data is presented as the mean ± standard deviation.
